# Distribution patterns of aquatic birds in a high-Andean wetland in southeastern Peru: an approach based on environmental factors

**DOI:** 10.1101/2025.03.01.640977

**Authors:** Carlos Lazo, Renny Daniel Diaz, Alexis Díaz, Victor Bustinza, Wilfredo Chavez, Aracely Dayana Machaca, Yoana Yarasca, Edwin Calderon, Maria Masias

**Affiliations:** Dirección de Investigación en Ecosistemas de Montaña, Instituto Nacional de Investigación en Glaciares y Ecosistemas de Montaña, Cusco, Cusco, Perú; Círculo de Investigación de Ornitología, Universidad Nacional Agraria La Molina, Lima, Lima, Perú; Centro de Investigación Vertebrate, Universidad Nacional San Antonio Abad del Cusco, Cusco, Cusco, Perú; División de Ornitología, Centro de Ornitología y Biodiversidad, Lima, Lima, Perú

**Keywords:** High-Andean wetlands, aquatic birds, distribution, environmental variables, Lake Piuray, Cusco, Peru

## Abstract

High-Andean wetlands play a crucial role in the conservation of avian biodiversity, acting as ecological oases amidst the arid Andes. However, these ecosystems face increasing anthropogenic and climatic pressures, jeopardizing both the ecosystem services they provide and the biodiversity they harbor. Moreover, research on these ecosystems remains scarce, limiting the knowledge necessary for their effective conservation. This study, conducted in Lake Piuray in the southeastern Andes of Peru, investigated the spatial and temporal distribution patterns of aquatic birds, assessing the influence of environmental variables such as depth and chlorophyll-a content. From December 2022 to November 2023, monthly surveys were carried out at 13 counting points distributed across four zones of the lake, encompassing natural and altered habitats such as lakeshore beaches and cultivated areas. Distribution maps were generated using weighted interpolation models and additive models to analyze non-linear relationships between bird abundance and environmental variables. Additionally, differences between seasons (wet and dry) and the evaluated zones were analyzed using non-parametric tests. A total of 43 aquatic bird species were recorded, with 19,768 individuals. Areas with low depth (<15 m) and intermediate to high levels of chlorophyll-a (I543 index: 0.20-0.25) concentrated the highest abundance and species richness. Zones with lakeshore beaches showed greater abundance and richness, while deeper zones exhibited the lowest values. At the family level, shorebirds preferred shallow waters, and diving birds tolerated greater depths. Although no significant differences in richness, abundance, and diversity were found between seasons at the community level, differences were identified in six families, with variations in their abundance between the wet and dry seasons. These results highlight the importance of shallow areas with high chlorophyll-a concentrations for the conservation of aquatic birds in high-Andean wetlands. This study is one of the few to analyze the influence of environmental and temporal factors on the distribution of high-Andean aquatic birds, and its representative nature makes it a valuable model for identifying priority areas for the conservation of these species.

## Introduction

The tropical Andes are one of the most biodiverse regions globally and represent a conservation priority [1]. However, these ecosystems face constant anthropogenic pressures (agriculture, mining, and overgrazing), particularly due to changes in land use and land cover [2,3]. In this context, wetlands located in the central Andes above 3500 m asl [4] stand out as critical conservation areas, as they are the primary providers of ecosystem services due to their high primary productivity and water regulation functions [5]. Additionally, these wetlands act as ecological “oases” because of their significant differences in productivity and biodiversity compared to the surrounding areas, especially during the dry season [6,7]. However, significant knowledge gaps remain regarding the extent, characterization, and biological richness of these wetlands in South America [8].

In the Neotropical region, wetland conservation and biodiversity are important research topics; however, many organizations focus their efforts on specific ecosystems, such as coastal wetlands and mangroves [9]. This trend is particularly evident in Peru, where most wetland studies have been conducted in coastal areas, with a strong emphasis on the wetlands of the central coast. This research imbalance contrasts with the limited attention given to Andean regions, despite the fact that the country hosts a similar proportion of coastal (46.6%) and Andean (47.2%) wetlands [10]. This highlights the need to expand research on Andean wetlands, particularly regarding biodiversity conservation [11].

Bird monitoring provides essential data for detecting environmental changes [12]. Birds, due to their sensitivity to anthropogenic disturbances, serve as key ecological indicators, especially in wetlands, where hydrological changes can significantly reshape their communities, leading to the decline of certain species and temporary increases in others [12, 14]. One example of this dynamic is the Lake Piuray, a wetland located in the tropical Andes of southeastern Peru, which, along with its source basin, is home to 147 bird species, including migratory, endemic, and some threatened species [15]. However, this lake faces increasing anthropogenic pressures, such as land-use changes and urban development, as well as climatic pressures due to variations in precipitation and temperature. As a result, a progressive decline in avian species diversity in Lake Piuray has been observed since 1993 [17].

The aquatic birds of Lake Piuray are particularly vulnerable, as the presence and abundance of these species in high-Andean lakes are closely dependent on environmental factors such as water depth and nutrient availability [18]. The distribution patterns of these species and their relationship with the different habitats of the lake remain unknown, which is crucial information for identifying priority areas and habitats in conservation efforts [19]. Therefore, the goal of this research was to determine the spatial-temporal distribution patterns of the aquatic bird community and families in Lake Piuray and to describe the environmental characteristics associated with these patterns (water depth and primary production). Additionally, the influence of the wet and dry seasons, as well as four distinct zones within the lake, on the richness, abundance, and diversity of aquatic birds was analyzed at both the community and family levels. This study is expected to contribute to the understanding of the factors influencing the distribution of aquatic birds, helping to identify key sites for the conservation of these species. Furthermore, the findings aim to be representative of wetlands in the central-southern region of the western Peruvian Andes, where knowledge about the factors affecting the distribution of aquatic birds remains limited.

## Methods

### 2.1. Study Area

Lake Piuray (13°25’0.00"S, 72°1’58.00"W; 2669 m asl) is located in the district of Chinchero, province of Urubamba, department of Cusco, Peru. This lake covers an area of 356 ha and reaches a maximum depth of 43.85 m. It is considered one of the main water sources for the city of Cusco, supplying 38% of its population [20]. The lake primarily receives its water recharge from ecosystems located at the headwaters of the Piuray micro-watershed [21]. The lake’s surroundings include agricultural areas, forest plantations, and urban zones, as well as two natural ecosystems: ‘humid shrubland’ and ‘humid puna grassland’ [15, 22]. In 2023, the area experienced an average annual precipitation of 809.6 mm, with two well-defined climatic seasons: a wet season (November - April) with 691.8 mm and a dry season (May - October) with 117.8 mm.

### 2.2. Selection of Sampling Sites

To facilitate the evaluation of the area and capture the specific characteristics of each region of the wetland, the study area was divided into four zones (Fig. 1) covering the entire perimeter of the Lake Piuray [23]. Zone 1 (Z1), located to the east, includes extensive agricultural areas and shallow lakeshore beaches. Zone 2 (Z2), situated to the south, also encompasses agricultural lands and a narrow strip of shallow lakeshore beach; however, the eastern end is dominated by considerably deep waters. Zone 3 (Z3), to the west, is characterized by extensive agricultural areas that extend almost to the shoreline, leaving little space for lakeshore beaches; here, nearshore waters are shallow but increase rapidly in depth toward the center of the lake. Finally, Zone 4 (Z4), to the north, is characterized by its proximity to eucalyptus plantations and agricultural areas located at the edge of the lake, with almost no lakeshore beach and deep waters.

**Fig 1.**
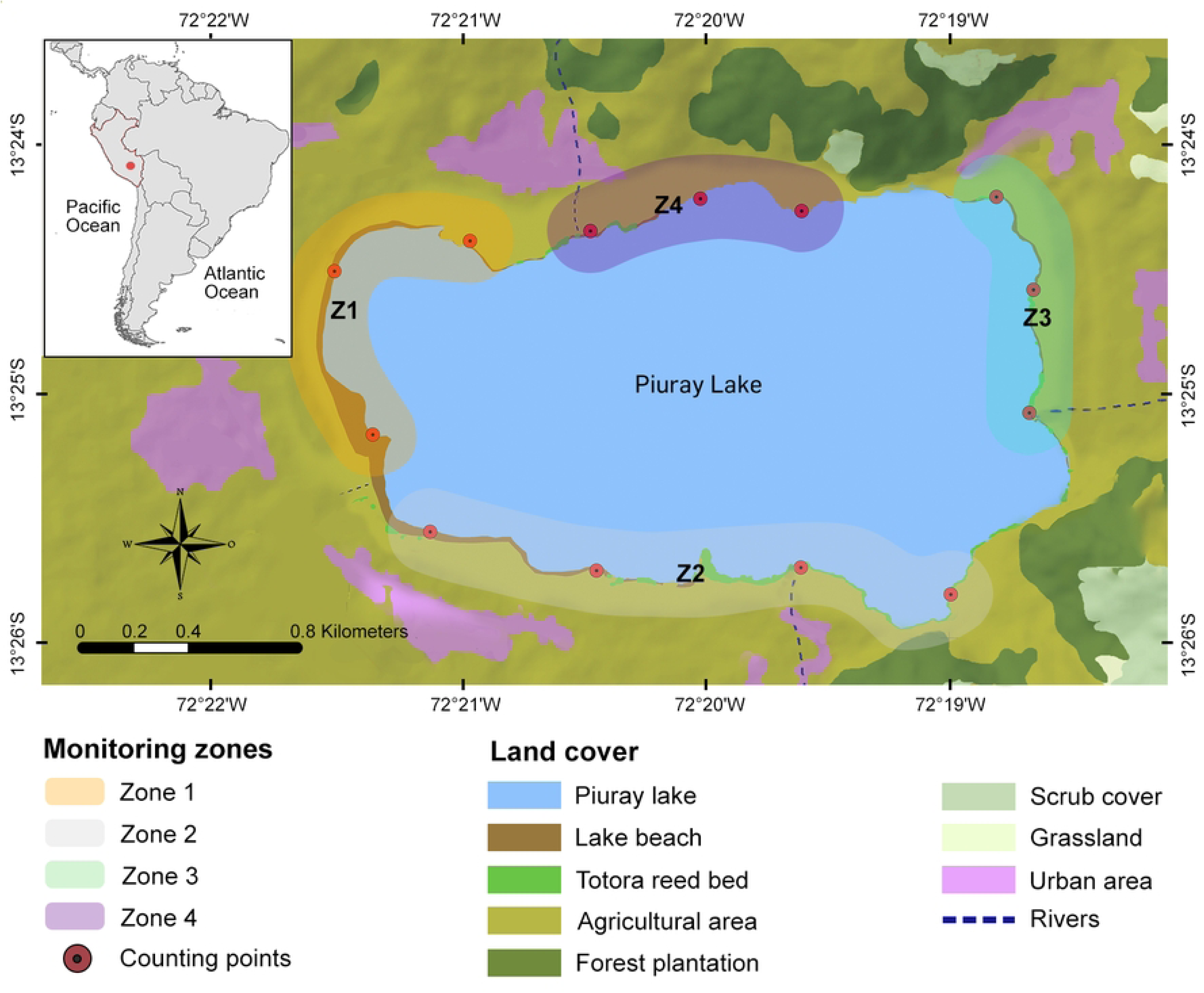
Location of study area and counting points on the land cover map of Lake Piuray and nearby areas. Land cover map obtained from INAIGEM [15]. https://doi.org/10.5281/zenodo.14902269

A total of 13 permanent counting points with a radius of 150 m were established [24]. Each of the four zones had three counting points, except for Zone 2, which had four due to its larger size. The average distance between points was 562.84 m (± 136.24 m standard deviation). Each counting point was located at the edge of the lake, completely surrounding it, in areas that facilitated the observation and identification of birds [25, 26].

### 2.3. Bird Censuses

Between December 2022 and November 2023, monthly bird censuses were conducted with four observer teams, one for each study zone. The censuses were carried out simultaneously to avoid double counting, following the protocol described by MINAM [25]. Each census began between 07:00 and 08:00 h and concluded between 10:00 and 11:00 h (PET). At each counting point, all birds seen or heard were recorded over a 15-minute period, with two consecutive repetitions within a 150 m radius. Observers used binoculars and cameras for bird identification and recorded data in field notebooks.

### 2.4. Data Analysis

A total of 11 families of aquatic birds were recorded in both seasons, with 34 species observed during the wet season and 39 species during the dry season. The analysis was conducted at the family level, as species of interest in this research, such as the Andean Ibis (*Theristicus branickii*) and the Buff-breasted Sandpiper (*Calidris subruficollis*), both threatened, as well as other migratory species of interest, had very few records during the study. Analyzing at the family level also allows for the identification of shared environmental response patterns for rarely observed species by combining data from multiple species with similar distributions [27].

#### 2.4.1. Determination of Abundance Patterns in the Aquatic Bird Community and Families

Two complementary analyses were performed: one at the community level and another at the aquatic bird family level. To visualize spatial abundance patterns in Lake Piuray and identify priority conservation areas, spatially continuous abundance distribution maps were generated using RStudio [28]. A Generalized Additive Model (GAM) was used, following the methodology proposed by Wood [29], to correlate the abundance of the community and each aquatic bird family (response variable) with chlorophyll-a content and lake depth (explanatory variables). A non-parametric model was chosen due to the expected nonlinear relationships between environmental variables and bird abundance [30].

To estimate the spatial distribution of birds in the study area, the Inverse Distance Weighting (IDW) interpolation method was applied. This method estimates abundance values in unsampled areas based on data from counting points, converting point data into continuous raster data for easier integration into spatial models. IDW is the most commonly used deterministic model in spatial interpolation [31, 32, 23]. Spatial interpolation methods assume a stronger correlation between nearby points than between more distant ones, a principle known as Tobler’s First Law of Geography [33]. IDW assumes that each measured point has a local influence that decreases with distance, meaning that closer points have similar values and greater influence on the interpolation, while more distant points are independent and have less influence [34]. In our study, the accumulated abundance values of the bird community and aquatic bird families at each counting point were used for interpolation.

Depth gradient data for the lake were provided by ANA, which conducted a bathymetric study, including a digital elevation model (DEM) [35]. To estimate chlorophyll-a content in Lake Piuray, we used the I543 index on Sentinel-2 satellite images, following the methodology described by Guigou [36]. Two study periods corresponding to the wet and dry seasons between 2022 and 2023 were selected. The images were processed using Google Earth Engine [37], where a customized script was implemented for data extraction and analysis. Image selection considered cloud-free periods for both study years. Before applying the I543 index, atmospheric and radiometric corrections were made to ensure accurate chlorophyll-a estimation.

To analyze the relationship between the distribution of the aquatic bird community and families and the physical variables of the lake (depth and chlorophyll-a), the ‘raster’ package [38] was used. The rasters generated by the IDW interpolation method were individually stacked with the DEM of Lake Piuray and the chlorophyll-a content raster. This resulted in a raster with a spatial resolution of 10 meters per pixel in each case. The GAM was then applied using the ‘mgcv’ package [39], chosen for its ability to capture complex nonlinear relationships between the response and explanatory variables without imposing parametric constraints [40]. A quasi-Poisson distribution with a logarithmic link function was used to address overdispersion in the data, and the Restricted Maximum Likelihood (REML) method was employed to fit the model. This approach allowed for flexible modeling of bird responses to environmental gradients, considering possible interactions between habitat structure (depth) and food availability (chlorophyll-a). The significance of each explanatory variable in the model was assessed with a threshold of P < 0.05. Additionally, the “gam.check” function in the ‘mgcv’ package was used to analyze and, if necessary, adjust the smoothing parameter (k) for each explanatory variable based on significance values.

#### 2.4.2. Analysis of Seasonal and Zonal Influence on the Aquatic Bird Community and Families

At the community level, species richness, diversity (Shannon-Wiener index), and total abundance (measured as the maximum cumulative count) were calculated for each season and in each of the four study zones. To evaluate differences between seasons and zones regarding species richness, diversity, and community abundance, as well as the abundance of aquatic bird families, the Kruskal-Wallis test was used, followed by post-hoc comparisons using the Dunn test with Bonferroni correction. These analyses were performed using the ‘FSA’ package [41] in RStudio, considering a significance threshold of P < 0.05.

## 3. Results

### 3.1. Abundance Patterns of the Aquatic Bird Community and Families

At the community level, the distribution maps obtained using the GAM model (Fig 2), based on IDW extrapolation, lake depth, and chlorophyll-a, showed similar explained deviance in both seasons: 44.1% for the wet season and 41.9% for the dry season. These maps reveal patterns with considerably high abundance in Z1 and the westernmost part of Z2, both in the wet and dry seasons, corresponding to the southeastern end of the lake. On the other hand, the lowest values were recorded in the central part of the eastern half, coinciding with the deepest areas of the lake (Z3 and Z4). The nonlinear relationships for the depth variable (Fig 2) indicate an inverse relationship between bird abundance and depth in both seasons, with a decrease in abundance at depths greater than 30 m and higher concentrations at depths less than 15 m. Regarding chlorophyll-a, the highest abundance values were recorded between 0.20 and 0.25 (chlorophyll index I543), while negative relationships were observed at values below 0.15.

**Fig 2.**
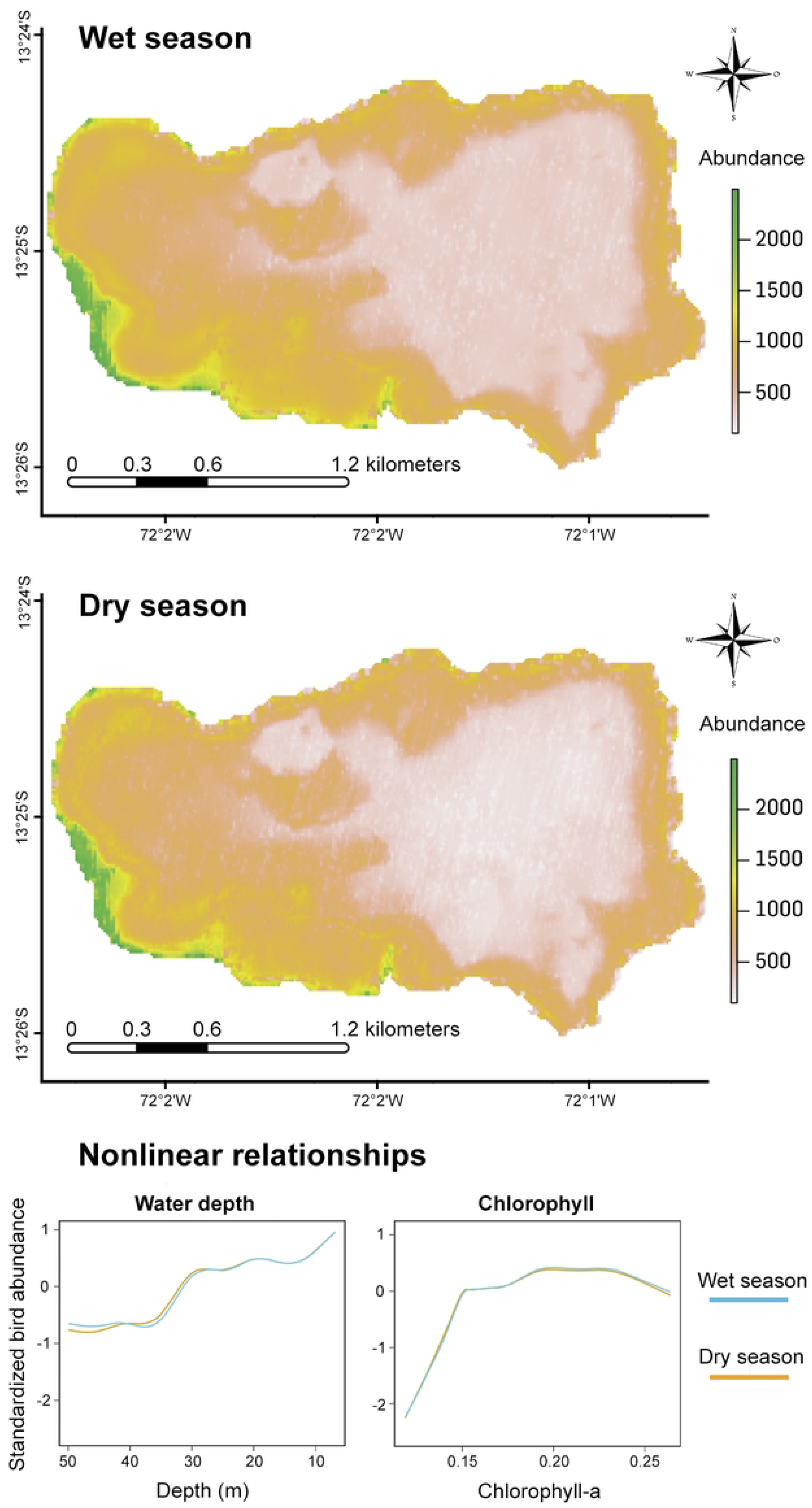
Distribution maps of the abundance of aquatic birds in Lake Piuray, estimated using a generalized additive model for the wet and dry seasons (above). The green shades represent areas with the highest concentration of individuals. Graphs of nonlinear relationships between environmental variables (water depth and chlorophyll-a concentration) and the abundance of aquatic birds in both seasons (below). Trend lines below zero on the Y-axis indicate negative relationships. https://doi.org/10.5281/zenodo.14902486

At the family level, the GAM model performed similarly for 9 out of the 11 families studied, with an average explained deviance of 35.9% (± 9.1%) in the wet season and 38.1% (± 4.4%) in the dry season. However, the families Ardeidae and Threskiornithidae showed significantly low values in both seasons. The adjusted distribution maps (Fig 3) reflected similar patterns for most families, with high concentrations between Z1 and the western end of Z2 and low concentrations in the deeper areas of the lake. However, the Threskiornithidae family showed an irregular distribution during the dry season. The families Charadriidae, Phoenicopteridae, and Recurvirostridae showed a more pronounced tendency for abundance in shallow waters compared to other families (Fig 4). In contrast, the Threskiornithidae family showed a relatively constant response in both seasons, suggesting a lower influence of depth on its abundance. Regarding chlorophyll-a, most families showed the highest abundance around 0.20 (I543 index), with negative relationships at values below 0.15, forming unimodal curves skewed toward areas with higher chlorophyll-a concentrations. However, the Threskiornithidae family exhibited a different response compared to the other families, as it did not show a defined peak or subsequent decline, indicating lower sensitivity to this variable compared to the other families.

**Fig. 3.**
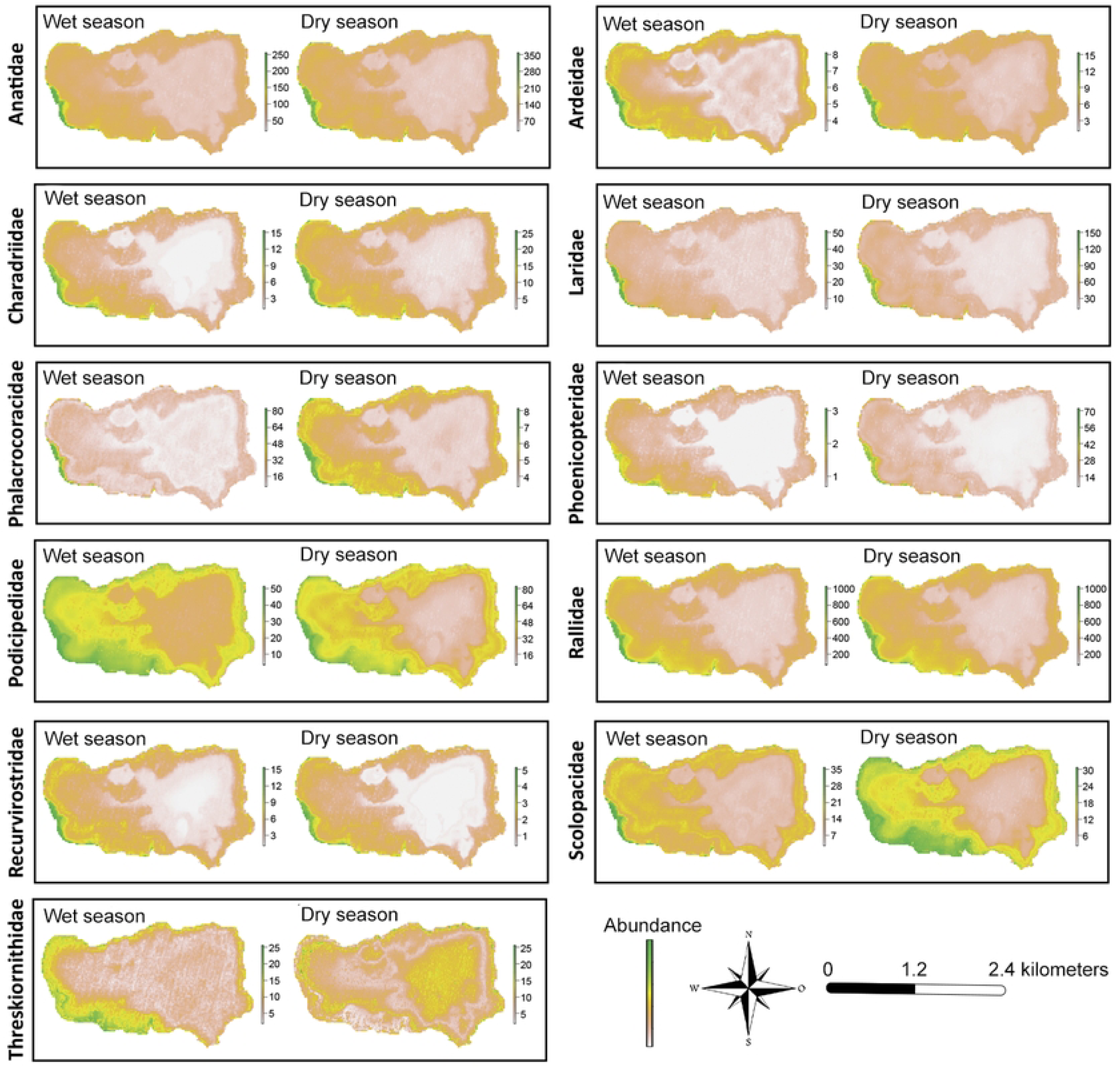
Distribution maps of the abundance of 11 aquatic bird families in Lake Piuray, estimated using a generalized additive model for the wet and dry seasons. Areas shaded in green represent zones with the highest concentration of individuals. https://doi.org/10.5281/zenodo.14902660

**Fig 4.**
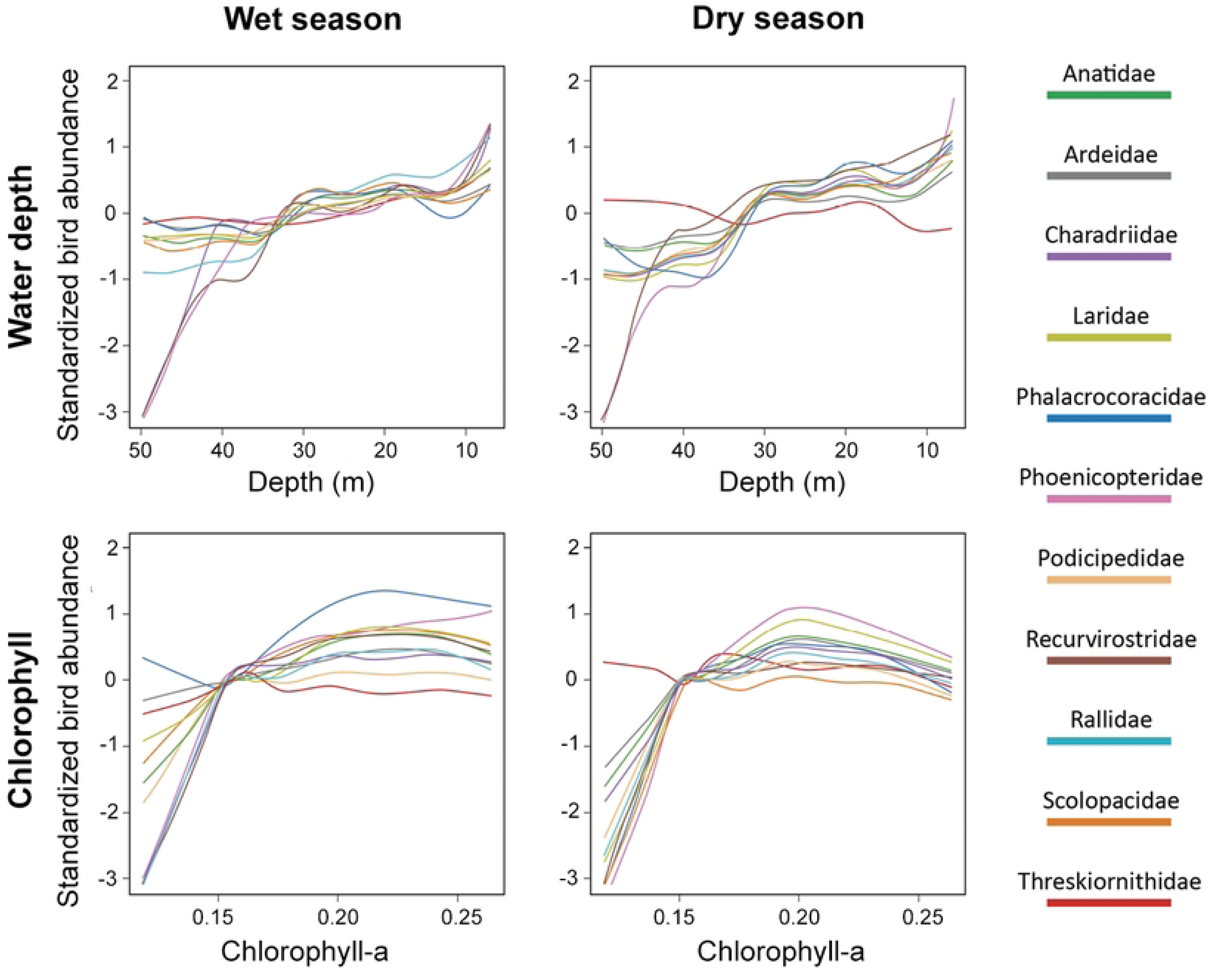
Nonlinear relationships between environmental variables (water depth and chlorophyll-a concentration) and the abundance of 11 aquatic bird families during the wet (left) and dry (right) seasons. The results were obtained using a generalized additive model. Each colored line in the graph represents a bird family, as shown in the legend. Trend lines below zero on the Y-axis indicate negative relationships, while lines above zero indicate positive relationships. https://doi.org/10.5281/zenodo.14902710

### 3.2 Influence of Seasons and Study Zones on the Aquatic Bird Community and Families

A total of 43 species of aquatic birds, distributed across 11 families, were recorded, with a count of 19,768 individuals over 12 monthly censuses conducted at 13 counting points and 4 transects used for the avian community analysis. Statistical analysis showed no significant differences in species richness, diversity, and bird abundance between the wet and dry seasons (P > 0.05). However, when evaluating the influence of different zones, significant differences were identified in species richness and abundance among the four studied zones (P < 0.01), although no differences were found in diversity (P = 0.9965). Post hoc analyses using Dunn’s test revealed significant comparisons (adjusted P < 0.01), indicating that Z1 had the highest species richness (37 species) and abundance (10,148 birds), in contrast to Z4, which showed the lowest species richness (24 species) and abundance (2,303 birds) (Fig 5).

**Fig 5.**
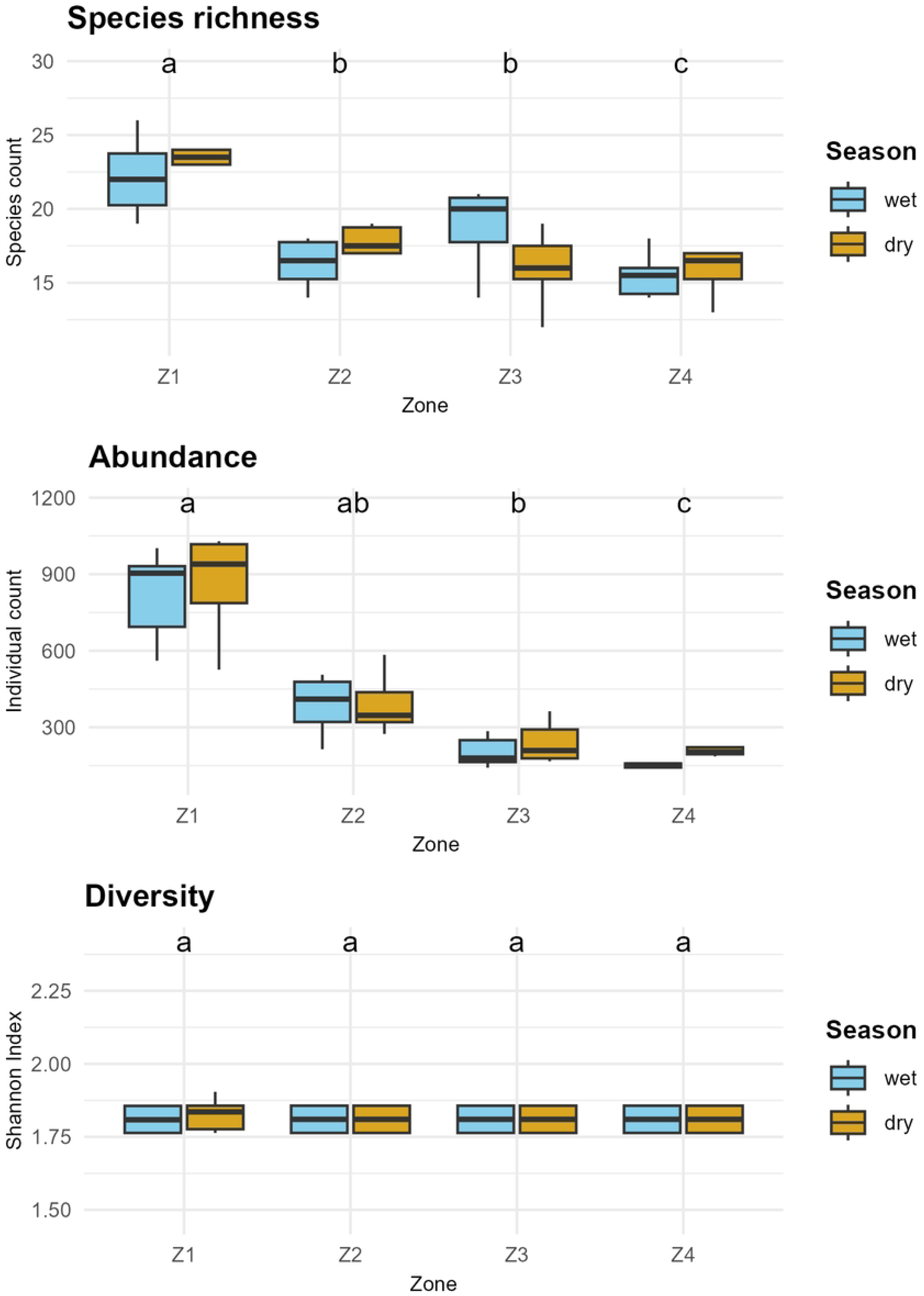
Comparison of species richness, abundance, and the Shannon diversity index between zones and seasons. The letters above the boxplots indicate significant differences between zones, as determined by the Dunn test with Bonferroni correction (p < 0.05). The colors represent the wet season (sky blue) and the dry season (dark yellow). https://doi.org/10.5281/zenodo.14902717

The family-level analysis revealed significant differences between the wet and dry seasons in 6 of the 11 analyzed families (P < 0.05). Specifically, the families Phalacrocoracidae, Scolopacidae, and Recurvirostridae exhibited higher abundance during the wet season, while the families Charadriidae, Laridae, and Phoenicopteridae were more abundant in the dry season. The remaining five families (Anatidae, Podicipedidae, Rallidae, Ardeidae, and Threskiornithidae) showed no significant differences between seasons (Fig 6). The comparison between zones for each family revealed highly significant differences in 9 of the 11 families (P < 0.01), except for Ardeidae and Threskiornithidae, which exhibited marginal significance (P = 0.04611) and non-significance (P = 0.07295), respectively. Post hoc analysis indicated that, for most families, Z1 showed significant differences compared to the other zones (adjusted P < 0.05), with higher abundance. In contrast, no significant differences were found in any comparisons for the Ardeidae and Threskiornithidae families (adjusted P > 0.05).

**Fig 6.**
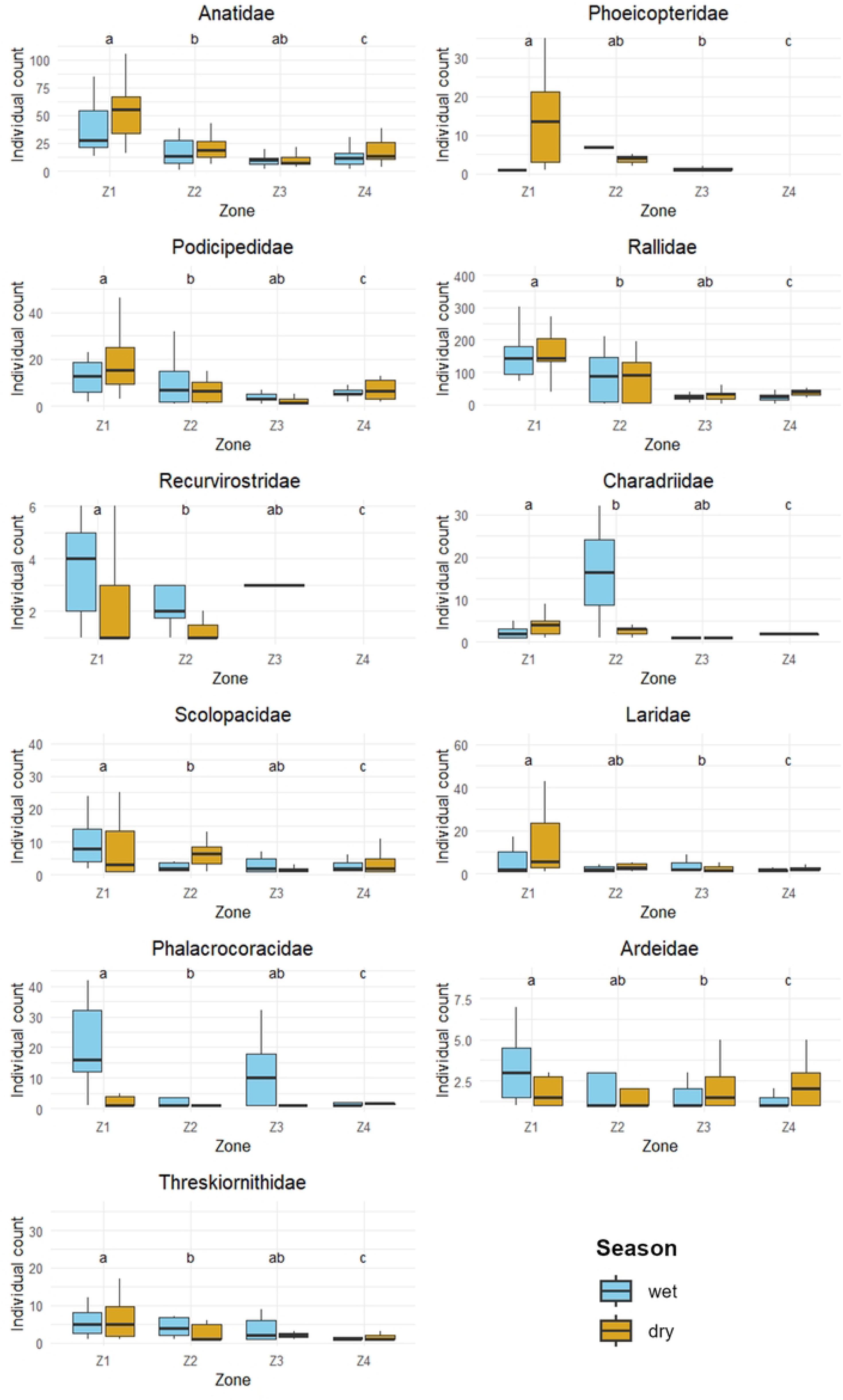
Comparison of the abundance of 11 families of aquatic birds across zones (Z1 to Z4) and seasons (wet and dry). The letters above the plots indicate significant differences between zones, as determined by the Dunn test with Bonferroni correction (p < 0.05). The colors represent the wet season (sky blue) and the dry season (dark yellow). https://doi.org/10.5281/zenodo.14902726

## 4. Discussion

This study represents the first analysis of the relationship between environmental factors and the distribution patterns of aquatic birds at two ecological scales (community and family levels) for a wetland located in the southeastern Peruvian Andes. The results make this work a representative contribution to understanding the wetlands of this region, which are characterized by a certain degree of biological similarity due to their shared geographic location, forming part of a network of high-Andean water bodies [42]. The richness of aquatic birds recorded in this study, with 42 species, is comparable to that documented in other wetlands of significant ecological importance in the central Andes, such as the Ramsar site of Lucre-Huacarpay, where previous research reported between 30 and 50 species [43,44]. Furthermore, this value is consistent with figures observed in larger lakes of the Andean region, such as Lake Junín, which hosts 55 species [45], or Lake Titicaca, with 45 species [46].

At the community level, the GAM model identified the highest concentrations of aquatic birds and positive nonlinear relationships in the shallowest areas of the lake (< 15 m) during both seasons. This pattern is consistent with previous findings in Andean lakes of Ecuador [18], in tropical coastal lakes in southeastern Brazil [47], and even in simulated scenarios modeling aquatic bird abundance as a function of depth [30]. These results support the importance of depth as a determining factor in the distribution and abundance of aquatic birds [48; 49]. Regarding primary production, represented by chlorophyll-a, the model showed that the highest concentrations of aquatic birds were associated with I543 index values between 0.20 and 0.26, corresponding to areas of higher productivity, consistent with other findings in wetlands of the eastern high Andes of Bolivia [50]. On the other hand, negative nonlinear relationships were observed at values below 0.15 in both seasons, reaffirming the strong correlation between this variable and the distribution and abundance of aquatic birds [51, 52]. Although Lake Piuray exhibits an initial mesotrophic state, characterized by a moderate amount of nutrients [53], the relationship between primary productivity and aquatic birds suggests that even moderate nutrient levels can significantly influence the dynamics of these communities.

At the family level, distribution patterns exhibited various trends, as factors such as depth can elicit differentiated responses among families depending on depth gradients [54]. For instance, shorebird families like Charadriidae and Recurvirostridae showed nonlinear relationships with peaks in abundance only in very shallow waters and highly negative responses in deeper waters, as these birds prefer extremely shallow waters (<0.10 m) [55]. Similarly, the Phoenicopteridae family, represented solely by *Phoenicopterus chilensis* in our study, preferred shallow waters with high primary productivity, a characteristic trait of this bird family [56]. Families with some diving species, such as Anatidae, Rallidae, and Podicipedidae, showed greater tolerance in moderately deep waters (20 to 30 m), as evidenced by distribution maps that place a significant number of individuals from these families in areas with such depths. This aligns with findings by Colwell and Taft [57], who describe a higher number of diving species in seasonal wetlands in the southern California valley. A different case was the Threskiornithidae family, composed of *Plegadis ridgwayi* and *Theristicus branickii*, which showed non-linear relationships close to zero for both depth and chlorophyll-a content across both seasons. Additionally, this family had the lowest explained deviation values of all, with 13.3% in the wet season and 4.08% in the dry season (Table 1). This suggests that depth and chlorophyll-a content are not variables that explain the distribution and abundance of this bird family, which is consistent with observations during monthly surveys, where the highest number of individuals from this family was seen in agricultural fields surrounding the lake, regardless of the water depth near the fields. The Ardeidae family also showed a low explained deviation, especially in the wet season (15.1%). The results for these last two families may be explained by the strong association that ibises and herons have with vegetation abundance rather than with other environmental variables [58].

**Table 1.** Model fit metrics for aquatic bird abundance across seasons.

|  | Wet season |  | Dry season |  |
| --- | --- | --- | --- | --- |
| Family | Adjusted R<sup>2</sup> | Explained Deviance (%) | Adjusted R<sup>2</sup> | Explained Deviance (%) |
| Anatidae | 0.394 | 38.9 | 0.333 | 36.4 |
| Ardeidae | 0.159 | 15.1 | 0.282 | 28.4 |
| Charadriidae | 0.368 | 45 | 0.331 | 40.8 |
| Laridae | 0.345 | 33.4 | 0.272 | 30.4 |
| Phalacrocoracidae | 0.238 | 20.8 | 0.376 | 40.7 |
| Phoenicopteridae | 0.309 | 44.1 | 0.328 | 42 |
| Podicipedidae | 0.32 | 30.7 | 0.328 | 32.5 |
| Recurvirostridae | 0.366 | 41.8 | 0.381 | 39.1 |
| Rallidae | 0.43 | 44.5 | 0.366 | 44 |
| Scolopacidae | 0.244 | 24.3 | 0.378 | 37.1 |
| Threskiornithidae | 0.153 | 13.3 | 0.0363 | 4.08 |

Adjusted coefficient of determination (adjusted $R^2$ ) and percentage of deviance explained by the generalized additive model (GAM) for the abundance of aquatic birds in 11 families during the wet and dry seasons. The values reflect the model's fit and explanatory power in relation to the environmental variables considered.
https://doi.org/10.5281/zenodo.14902748

Regarding temporal variations, at the community level, no significant differences were found in richness, abundance, and diversity between the wet and dry seasons. This result is consistent with previous research conducted in wetlands of the central and southern Andes of Peru [6, 59] and could be explained by the high presence of resident species, including predominant ones. On the other hand, spatial variations were observed according to the zonation at the community level. Zone Z1 stood out as the area with the highest bird richness and abundance, associated with the largest extent of lakeshore beaches compared to the other zones. This factor has also been key in explaining bird richness and abundance in other Andean lakes, such as Lake Tota in the Eastern Cordillera of the Colombian Andes [60]. However, diversity remained similar across all studied zones, possibly due to the predominance of species like *Fulica ardesiaca*, which maintains abundant populations in these ecosystems [18, 61].

At the family level, significant variations in the abundance of six bird families were identified between the dry and wet seasons. During the rainiest months of the wet season (December to February), the Phalacrocoracidae family, represented exclusively by *Phalacrocorax brasilianus*, and the Recurvirostridae family, dominated by *Himantopus mexicanus*, reached their peak abundances. This pattern is consistent with data from the eBird portal, which reports higher concentrations of these species in high-Andean lakes during these months [62]. On the other hand, the Scolopacidae family, comprising 11 species classified as boreal migrants [63], showed higher abundance during the wet season, although some individuals remained in the region during the dry season. This permanence could be explained by oversummering strategies, as reported in some shorebirds on the central coast of Peru [64]. During the dry season, the Charadriidae and Laridae families exhibited different abundance patterns: Charadriidae, with four species but dominated by *Vanellus resplendens*, and Laridae, with two species but predominantly *Chroicocephalus serranus*, reached their highest abundance levels in the driest months (June to August), although both families maintained considerable populations throughout the year. Notably, *Vanellus resplendens* was reported to exhibit population movements from the headwaters of the basin toward Lake Piuray during the driest months [15]. Additionally, *Chroicocephalus serranus* is known to form flocks in the southern Andes of Peru during the dry season [65]. For the Phoenicopteridae family, *P. chilensis* was the only representative and was observed only during the dry season and in November, with no records in other periods of the year. This constitutes one of the first documented records of its wandering movements in lake Piuray. The changes in the population of these six bird families, alternating between the dry and wet seasons, could be one of the reasons why no significant difference was found in the aquatic bird community between the two seasons. Notable differences were also observed among the different study zones, particularly in Zone Z1, which hosted the highest abundance for most families, except for the Ardeidae and Threskiornithidae families, which showed no significant differences between zones. Zone Z1 was characterized by having the largest areas of shallow waters and lakeshore beaches, factors closely related to high richness and abundance of aquatic birds [66]. In contrast, Zone Z4 had the largest extent of deep waters and an almost nonexistent lakeshore beach, in addition to being close to a rural road and a eucalyptus forest, which likely explains the low levels of richness and abundance in this zone.

The conservation of Andean wetlands requires spatially precise predictions that consider both natural and anthropogenic aspects to identify the most important sites for conservation efforts [67]. In this study, the distribution patterns of aquatic birds appear to result from the interplay between natural and semi-natural habitat factors present in all study zones. The GAM models, which analyzed the influence of depth and chlorophyll-a concentration on aquatic birds, explained between 35% and 44% of the observed variability in the distribution of these species at the community and family levels, with depth being the most influential variable in all cases. These results suggest that other factors may also play an important role in the distribution of birds in lake Piuray. These could include local environmental factors, such as the presence or absence of emergent vegetation, which influences species that use it as refuge [68], or the influence of dominant generalist species, which can displace less abundant species [18], as is the case with *Fulica ardesiaca*, which has a large population in the lake Piuray [15]. Additionally, other factors such as wetland size, which in most study cases shows a positive correlation with species richness [69], water chemistry parameters like pH, which is associated with certain bird guilds [70], or urban development, which tends to reduce bird richness in wetlands [71], could also be determinants. Therefore, it is essential to include a greater number of variables in future studies to gain a more comprehensive understanding of the factors explaining the distribution of aquatic birds in these ecosystems.

## 5. Conclusions

Our findings contribute to identifying priority areas for the management and conservation of aquatic birds in lake Piuray, a representative area of the humid puna wetlands. We found that the shallowest areas of the lake with considerable chlorophyll-a content host the highest number of individuals and species of aquatic birds at the community level and for most families in both seasons. Shorebird families showed a greater affinity for very shallow waters, while diving bird families exhibited greater tolerance to deeper waters. We also found that the Threskiornithidae family does not show distribution trends strongly associated with depth and chlorophyll-a. Overall, our observations underscore the relevance of shallow waters with high chlorophyll-a as areas of great importance for the conservation of aquatic birds.

## Acknowledgments

Abril Rado, Arturo Mamani, Billy Ttito, Claudia Ibarra, Daniela Sequeiros, Diego Gayona, Franck Cusi, Marianela Vargas, Israel Huaranca, Johan Saire, Leidyjean Llanos, Luis Uñunco, Shirley Sarango,Sofia Alvares, Susan Blanco, Rosely Sotomayor, Vanessa Barbachan, Xiomara Ampuero.

## Acknowledgments

We sincerely thank the Instituto Nacional de Investigación en Glaciares y Ecosistemas de Montaña (INAIGEM) for their support in the development of this research project. We also extend our gratitude to the Círculo de Investigación en Ornitología (CIO) of the Universidad Nacional Agraria La Molina and the Centro de Investigación Vertebrate of the Universidad Nacional San Antonio Abad del Cusco for their valuable assistance in providing volunteers for this project.

## Supporting information

**S1 File. List of aquatic bird species and their abundance by season in Lake Piuray**

**S2 File. Bird abundance data at the community and family levels for creating maps using IDW interpolation.**

**S3 File. Raster data of bathymetry and chlorophyll-a for the wet and dry seasons in Lake Piuray.**

**S4 File. Data for analyzing the influence of seasons and zones on the community and families of aquatic birds in Lake Piuray.**

**S1 Table. Results of the Kruskal-Wallis and Dunn test for the comparison of aquatic bird families across seasons**

**S2 Table. Results of the Kruskal-Wallis and Dunn test for the comparison between zones by families**

## Notes

### Competing Interest Statement

The authors have declared no competing interest.

